# Anthropocentrism as a source of sampling bias in the fossil record

**DOI:** 10.64898/2026.03.04.709576

**Authors:** T.I.F. Foister, O.E. Wilson

**Affiliations:** Department of Geosciences and Geography, University of Helsinki

**Keywords:** fossil prospecting, collection bias, humans, anthropocentrism, spatiotemporal variation

## Abstract

The fossil record is the most important tool in palaeosciences, so continually reviewing and attempting to reduce biases in its collection is necessary to curate the best possible record of past life on Earth. Biases in the fossil record are introduced through both biological processes and data collection. Here we have investigated the extent to which anthropocentric data collection has contributed to sampling bias in the assembly of the current fossil record. We have found that the current fossil record (represented in this study by the NOW database) is anthropocentrically biased, both temporally and spatially. Specifically, fossil locality density is higher in time periods when hominins are found, and in known hominin-bearing locations. This demonstrates the need to stop essentializing the narrative of human evolution in paleoscience to reduce bias in sampling of fossil localities.

## 1. Introduction

The fossil record is the defining tool of palaeosciences as the only direct evidence of the geographic and temporal distribution of traits, species and lineages. The fossil record preserves only a very small percentage of all organisms which have lived on Earth, and the chances of any particular organism being preserved are determined by a combination of factors (e.g. depositional environment, taxonomic preservation potential and body size) (Behrensmeyer et al., 2000, 2003; Walker et al., 2020). As a result of this, the fossil record is full of numerous biases. Some of these biases are a function of the fossilisation process and cannot be removed, whereas human-introduced biases can be mitigated (Behrensmeyer et al., 2000; Nanglu & Cullen, 2023).

Sampling biases, which relate to how likely a fossil is to be collected and recorded in the fossil record, are one example. Within the mammal fossil record, large taxa have traditionally been of more interest, so they are more represented in the fossil record, while micromammals have been neglected (Bush et al., 2007; Cooper et al., 2006; Whitaker & Kimmig, 2020). Here we consider how focusing on a single taxonomic group (in this case hominins) can be considered as an example of sampling bias on a global Cenozoic scale.

Anthropocentrism is a complex concept used in many disciplines with varied meaning. Here, we use it to refer to the centering of *Homo sapiens*, its ancestors and related species, in the narrative of evolution. In our analysis we have selected all species in the tribe Hominini, and as such define anthropocentrism as the focus on the taxa within Hominini. In primatology, anthropocentric bias is clearly identified where primates are primarily studied with the goal of using them as a proxy to study human evolution. Consequently, non-human apes are more studied than non-hominoid primates, as they are closer relatives of humans (Ramsay & Teichroeb, 2019; Sayers, 2014). Many of these studies focus primarily on social and cognitive behaviours of interest to human evolution (Boyd, 2017; Radhakrishna & Jamieson, 2018; Ramsay & Teichroeb, 2019).

One of the critical questions facing all scientific disciplines today is inescapably the current biodiversity crisis. It has been argued that the most valuable contribution palaeosciences can have is through providing a baseline of undisturbed communities and ecosystems (Kiessling et al., 2019; Tierney et al., 2020). Moreover, if we wish to understand how ecosystems will respond to climate change and to prepare for and limit climate change and biodiversity loss we must look to pre-hominin conditions in deep time (Barnosky et al., 2017; Boivin et al., 2016; Jones et al., 2021), e.g. the Miocene (Steinthorsdottir et al., 2021) and Eocene (Burke et al., 2018), as it has been hypothesised that post-climate change conditions will most resemble those experienced in these time periods (Tierney et al., 2020).

This is not however, to say that investigating human evolution is without its merits, even within the context of climate change. In the long-term, investigating hominin evolution with the goal of understanding the changing human-nature relationship may provide insight to how anthropogenic forcing on the climate has shifted through time, and how this can be mitigated. Additionally, it is logical that as humans we are interested in the origins of our own species and how we came to exist in our present form. Research into this is valuable and has contributed to many important advancements both culturally and scientifically. For example, human origins research has been integral to dismantling institutionalised racism and contributed to ‘post-colonial’ theory (Porr & Matthews, 2017). Research into human origins intersects with many other fields and provides insights important to developing our understanding of the human condition. For example, human origins models have contributed to our understanding of human metabolism and thermoregulation by providing insight to how and why humans adapted to control their internal temperature, and how this is affected by traits such as bipedalism (Haman & Blondin, 2017). Whilst these contributions and human origins research as a whole remain valuable, by essentializing the narrative of human evolution in palaeoscience the data available to answer other questions become limited and biased, thereby restricting the impacts palaeosciences may have. For example, megafaunal extinctions have long been viewed through an anthropogenic lens, with a heavy focus on implications of how the human fossil record can be applied to interpret general palaeobiological processes (Amir et al., 2022; Braje & Erlandson, 2013; van der Kaars et al., 2017).

In this paper we have used the NOW database (The NOW Community, 2024) as a representative database of fossil species and localities to test our hypothesis that the fossil record is biased in favour of time slices and locations where hominins are known.

## 2. Materials & Methods

### 2.1. Data

In order to investigate if the fossil record is biassed we used the NOW database (The NOW Community, 2024) to assess the density of sites recorded in space and time. We exported the NOW database on 31-03-2023. We identified which locations contained hominin fossils (taxa within the subtribe Hominina and genus *Homo*), then plotted all locations along with the density of hominin sites (Fig. 2). Due to the nature of the NOW database, in this analysis, hominin sites means sites bearing hominin fossils, so that sites only with lithics or other anthropogenic material are not included. We also conducted this analysis at three time slices, the Cenozoic (all locations in NOW), Pliocene and Quaternary (Max Age < 5.3 million years ago) and Quaternary (Max Age < 2.6 million years ago, after the appearance of *Homo* genus). To specifically analyse the changes in density of sites in space, we tested if there is a greater number of sites in countries where hominini have been recorded. Hominin density was calculated using a geospatial kernel density estimate through the *eks* package (Duong, 2023) for R version 4.1.0 (R Core Team, 2021). All figures were produced using *ggplot2* (Wickham, 2016).

To investigate how the density of sites in NOW changes across time, we sorted the exported data into time bins instead of location. We visualised these data in two ways to interpret the density. First, we plotted a histogram of the number of sites across time, colouring the bars by epoch. Secondly, we plotted the count of sites by epoch, standardising for the length of epoch (number of sites/length of epoch (millions of years, (Ma))).

We used the Sedimentary Basins dataset (Nyberg & Howell, 2015a) from the Eco-ISEA3H database (Mechenich, 2022; Mechenich & Žliobaitė, 2023) for centroids positioned at a distance of ∼50 km (resolution 9 in the Eco-ISEA3H database) to identify the position of any modern sedimentary basins. The original dataset from Nyberg and Howell, (2015b) used to generate the Eco-ISEA3H version mapped the modern terrestrial sedimentary basins to identify the patterns of bias that exist in terrestrial basins today and identified whether these patterns might be applicable to the terrestrial fossil record. We use modern basin coverage as a proxy for sediment availability in the Cenozoic and overlaid the NOW localities to visualise any correlation, with sediment availability defined as the probability of encountering a sedimentary basin at each centroid of each of the hexagons defined in the Eco-ISEA3H database. Clearly, this modern coverage differs from the distribution of sedimentary basins in the fossil records, but we were unable to find any data that would allow us to compare basin availability in different periods of geological time. In addition, we note that there are patterns in basin distribution in the modern world that are likely to have been true in the past too, for example most modern basins are in arid environments, whilst snow or ice-covered environments are typically underrepresented (Nyberg & Howell, 2015b). We therefore use it as a proxy for where potentially fossiliferous sedimentary basins are likely to have been deposited, but acknowledge the challenge in this approach. This variable does not account for the accessibility of the sedimentary basin, or possible post-depositional effects. As such, it is only an indicator of where potentially fossiliferous sedimentary basins are likely to have been deposited.

### 2.2. The NOW Database

This work relies upon the NOW (New and Old Worlds) database as a proxy for the fossil record of the Cenozoic. The NOW fossil mammal database was first publicly released in 1996, going online in 2005. Originally, the database focused on the European Miocene and Pliocene (e.g. Fortelius et al., 2002), giving rise to its original title “Neogene Old World” but it has since expanded to be a global Cenozoic land mammal record, recording taxonomic, dating and locality information (Žliobaitė et al., 2023). The NOW database results from the work of a large community of collaborators and continues to be an actively expanding and changing database of fossil mammals. A detailed explanation of how these data are curated can be found on the database homepage (https://nowdatabase.org/). The NOW wiki (https://github.com/nowcommunity/NOW-Django/wiki) provides a detailed account of all data fields within the database. While the NOW community continually updates and curates the database there will always be inherent bias in the data contained within it, as for all databases based on the literature. For example, while attempts are constantly made to include data from publications that are not in English, there is likely to be a bias towards English publications due to the nature of publication (Amano & Sutherland, 2013a; Haddaway et al., 2020). Similarly, while attempts are made by the curators to include historical manuscripts as far as possible, there may also be a bias towards younger publications, a phenomenon which has been described for other fields as “citation amnesia” (Singh et al., 2023) or the “Ageism of knowledge” (Gottlieb, 2003). Such challenges are present for all databases built on literature searches, and we found the NOW database to be the best available proxy for the fossil record, as have other authors (Prothero, 2015; Žliobaitė & Fortelius, 2022). At the time of download (31-03-2023) the NOW database contained 7177 localities, 16322 species, and 75585 locality-species that are public entries. See Figure 1 for an example locality record. While NOW does not cover all fossil localities and specimens of the Cenozoic, partly as a consequence of its origins, it serves as an excellent representation of the mammalian fossil record, and given the attention given to updating taxonomy and chronology, is likely more accurate for mammals than other alternatives like the Paleobiology Database (PBDB) (Peters & McClennen, 2016; Žliobaitė & Fortelius, 2022). Moreover, as the NOW database originated as a Neogene palaeontological database we can ensure that the criteria and procedure for data entry have not been biased by an anthropology or archaeology focus, as other resources may have been. While the database can be applied to palaeoanthropological research, it is first and foremost a paleontological resource, and as such any anthropocentric bias discovered is a function of fossil prospecting/collection bias, and not of the database itself. On the other hand, this does mean that many sites in the Holocene which only have archaeological material are not captured in the NOW database, as they do not have faunal data to record. This means that our analyses may in fact underestimate the sampling bias generated by anthropocentrism, as human sites are more abundant in the Holocene.

**Figure 1.**
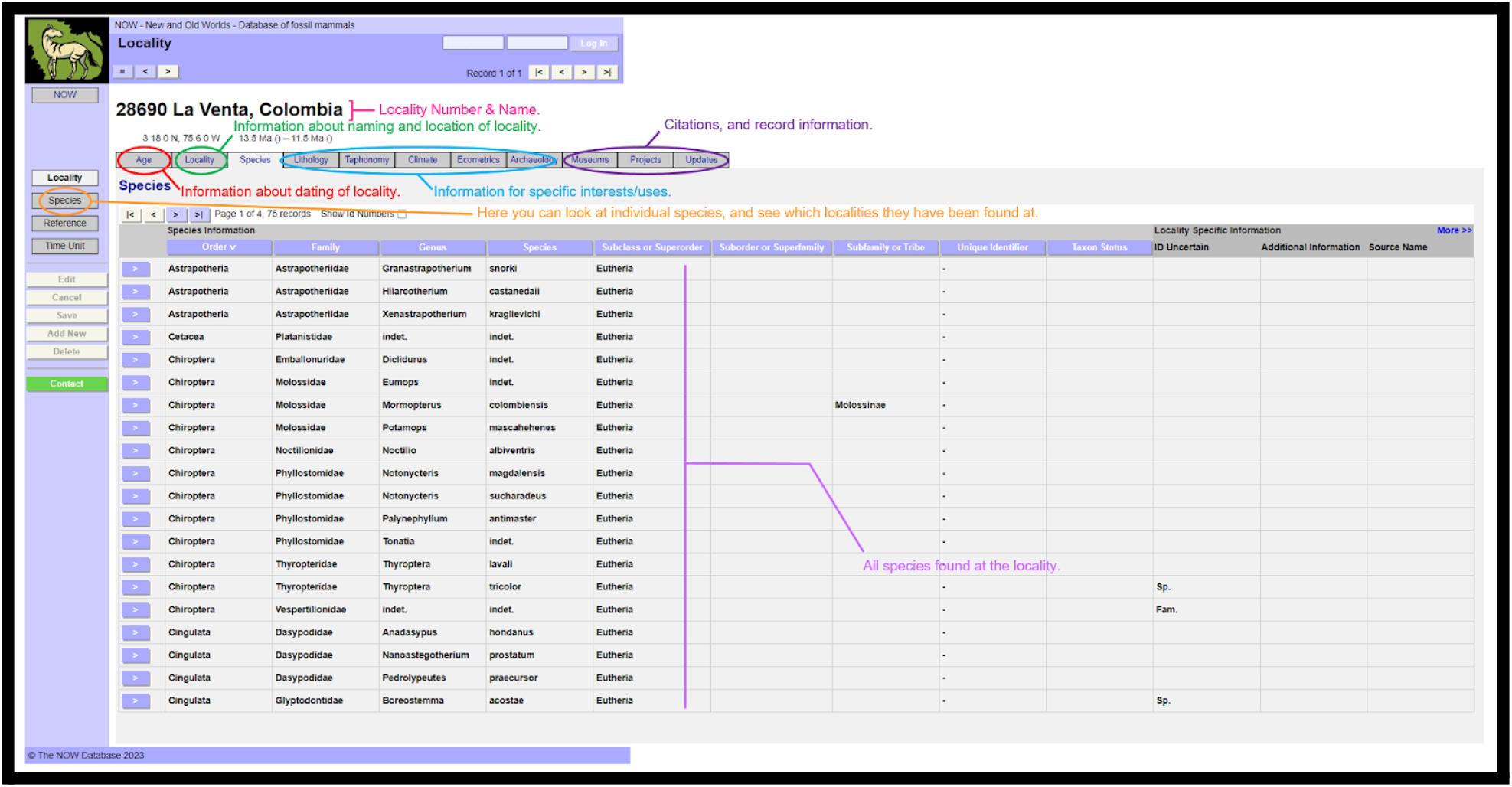
An example record from the NOW database, showing what information is available in the database for the Middle Miocene locality of La Venta, Colombia. This screenshot was taken at 14:57, on 12-06-2024.

**Figure 2.**
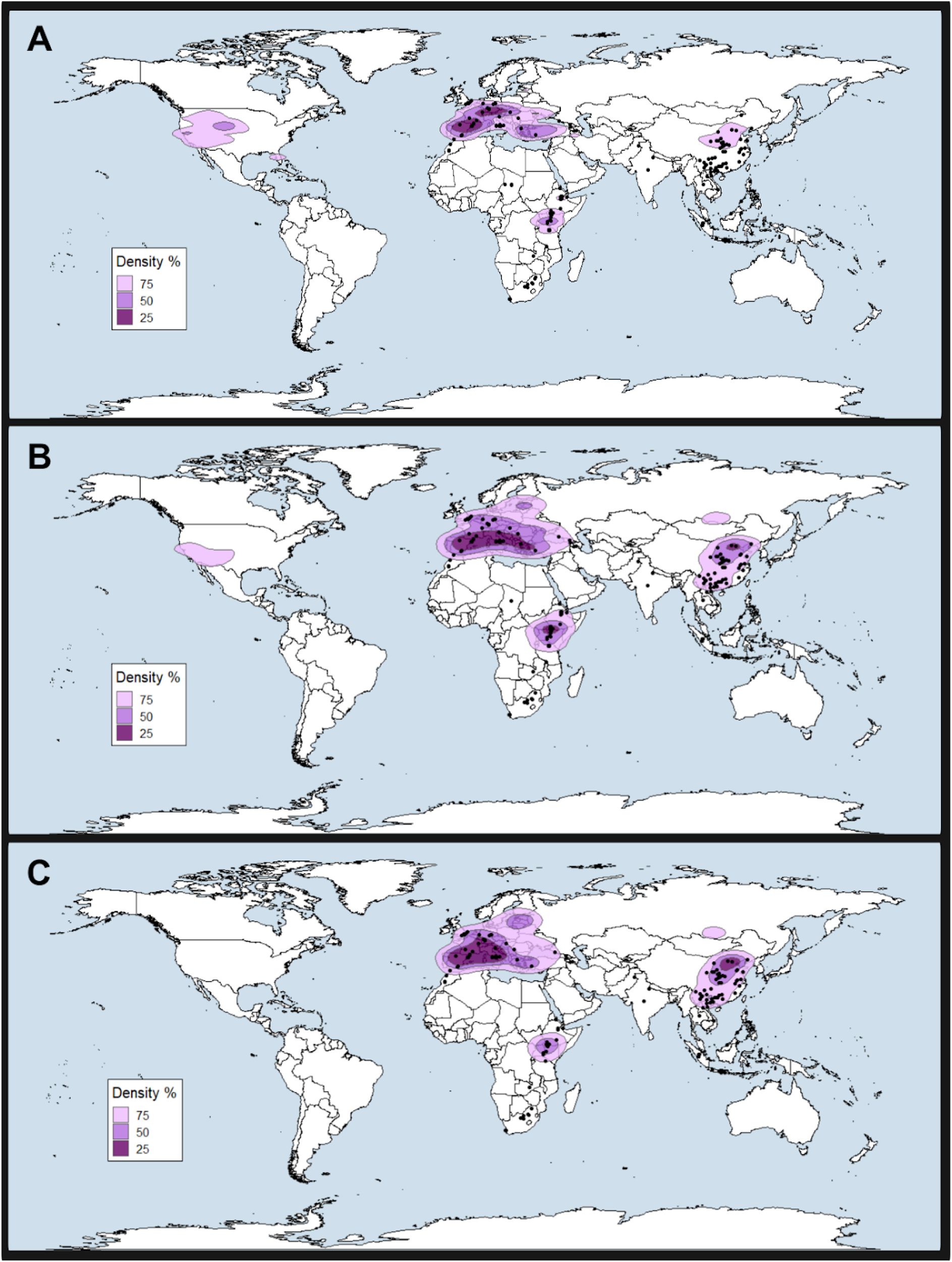
Density of fossil mammal localities from different time periods (purple polygons) and records of hominins from each period. Both densities and hominin occurrences taken from the NOW database of fossil mammals. A) Cenozoic B) Quaternary & Pliocene C) Quaternary.

## 3. Results

### 3.1. Spatial bias

Our results show that regions where, and time slices when hominins are found are highly sampled, while regions and time slices without hominins are relatively undersampled. The densest sampling is seen in regions containing hominin localities, such as Spain, eastern Africa and China (Fig. 2). The USA does not fit this pattern, with a high density of fossil localities where there are no hominin fossils (Fig. 2A, Fig. 3). However, these sites are mostly pre-Pliocene, and therefore predate the evolution of hominins. In addition, many of the oldest North American records in NOW result largely from the input of specific literature (e.g. Janis et al., 1998), so therefore have a history separate from the rest of the NOW database. The density pattern is strongest in Quaternary fossil localities, after the appearance of the *Homo* genus (Fig. 2C), with Quaternary localities especially concentrated in countries which contain hominin fossils. These analyses suggest that the Quaternary fossil record is more spatially anthropocentrically biassed than the Cenozoic as a whole.

**Figure 3.**
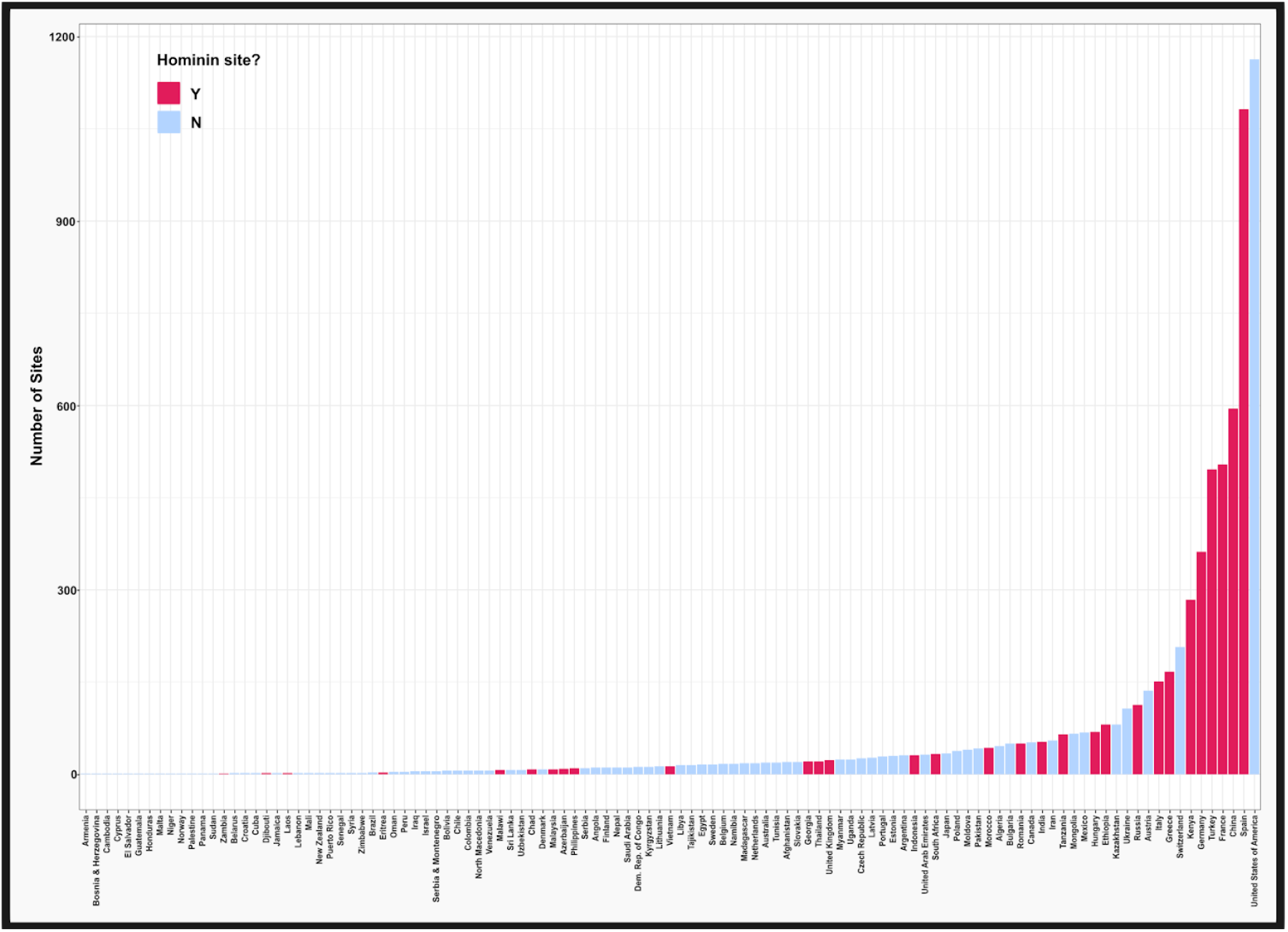
Records of fossil mammal localities recorded in the NOW database by country for the whole Cenozoic. Countries included here are all those with at least one locality recorded in the database. Countries are coloured by the presence or absence of any hominin fossils.

We identify South America, Australia and Western Africa as severely undersampled in the NOW database. Additional undersampled regions are Northern Eurasia (e.g. the Nordics, Russia, Kazakhstan and Mongolia) and Northern Africa (e.g. Egypt, Libya, Algeria). This indicates that the fossil record is biased in favour of sampling hominin locations. The distribution of sedimentary basins does not match the distribution of fossil localities (Fig. 4). Widespread sedimentary basins are present in many of these undersampled regions (though to a lesser extent in northern Eurasia), suggesting that the lack of localities in these regions is not solely due to a lack of fossiliferous sediment.

**Figure 4.**
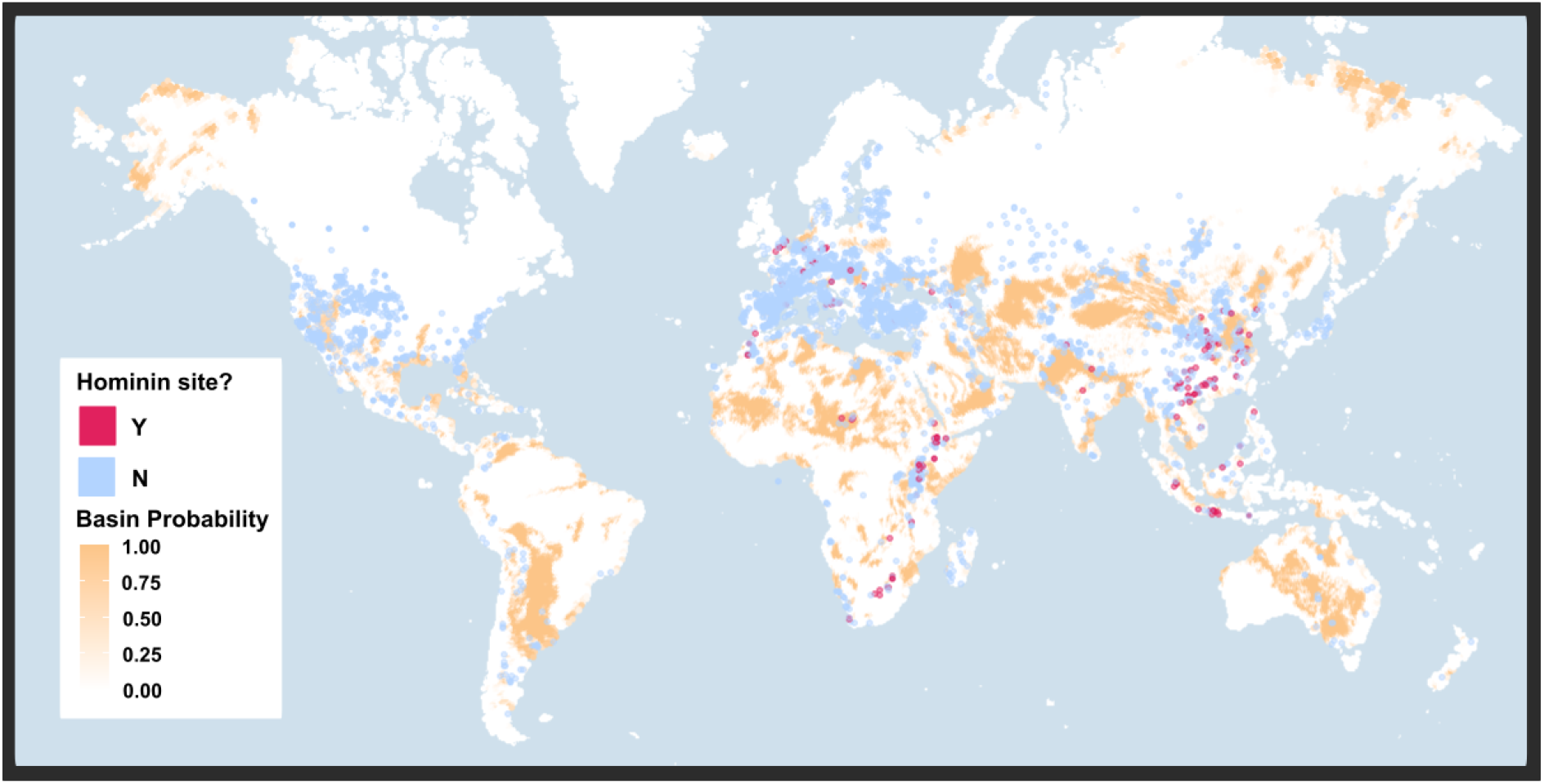
Global map of sediment availability using Sedimentary Basins dataset of the Eco-ISEA3H database. Showing probability of encountering a sedimentary basin at centroid (orange), with the distribution of fossil localities (from NOW database) overlaid. Fossil localities are coloured by the presence or absence of hominin fossils.

### 3.2. Temporal bias

The Miocene has the highest number of sites by a significant proportion, followed by the Pleistocene and Pliocene (Fig. 5). We expect the Miocene to have a high site number in the NOW database due to both the origins of the database(Žliobaitė et al., 2023), and the length of the epoch. When we standardise the number of sites to factor in the time length of the epochs, we see that the coverage of the Holocene is much higher compared to the other epochs, followed by the Pleistocene and Pliocene. This is presumably a result of both the shorter period of time represented and the sampling. The temporal coverage of fossil localities increases as we move through hominin evolution, with post-*Homo* time slices having the highest coverage (Fig. 5). Our analyses indicate that non-hominin-bearing time slices and regions are undersampled. This undersampling increases with time from hominin evolution, so that the Paleocene is the least sampled epoch, followed by the Eocene and Oligocene, though we acknowledge the length of these epochs relative to the Holocene, Pleistocene and Pliocene.

**Figure 5.**
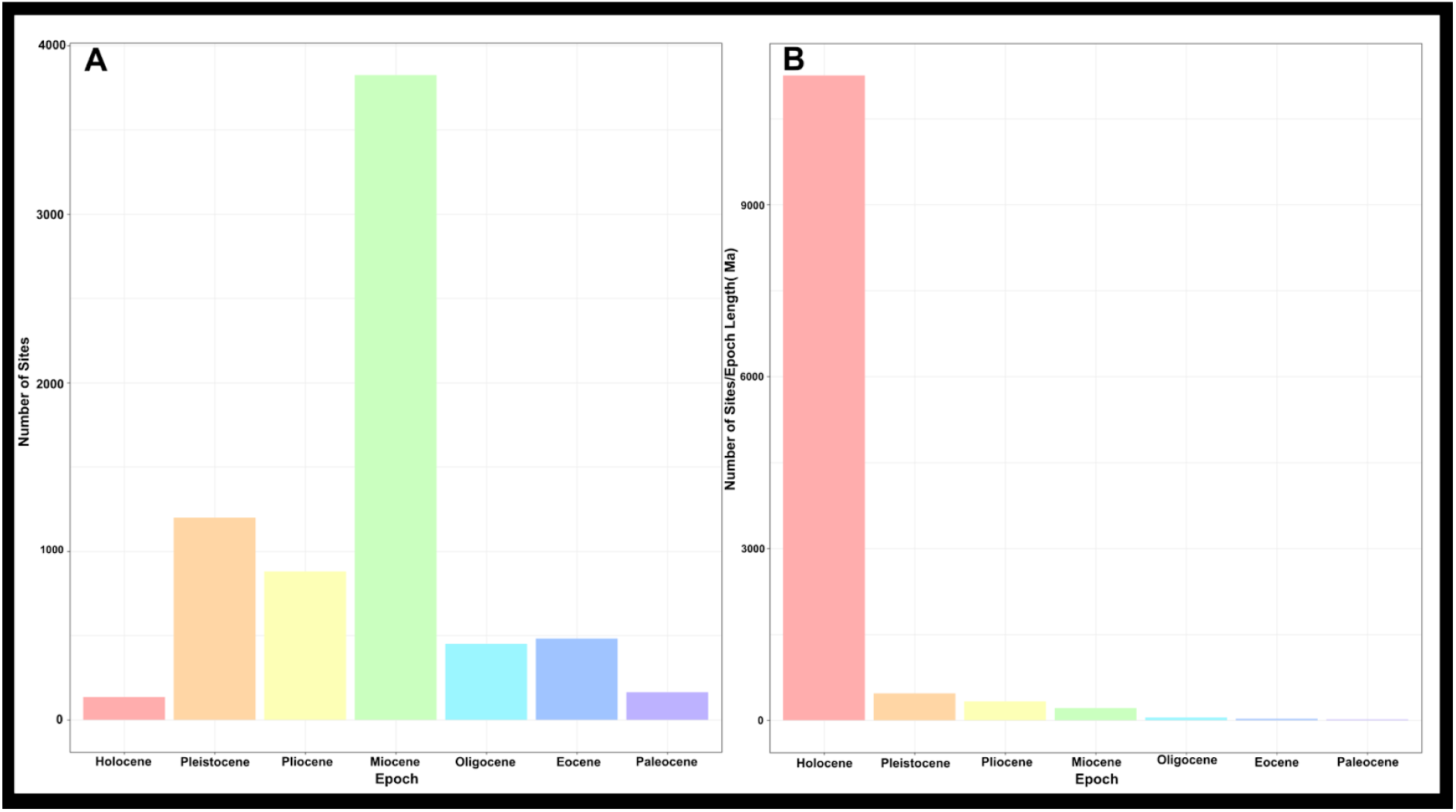
Bar plot showing the number of fossil mammal localities (from the NOW database) per epoch A) Raw count of fossil localities B) Number of fossil mammal localities standardised for length of epoch

## 4. Discussion

Our results indicate that anthropocentrism in fossil sampling can be considered as a source of bias in the fossil record. This bias is present temporally and spatially, as hominin bearing locations and time slices are better sampled compared to non-hominin bearing locations and time slices. In particular Australia, South America and Western Africa are undersampled, as shown in Figures 2 and 3, whilst this may be exacerbated by a lack of integration in the database (for example, the database is actively growing its South American data), it is also a reflection of a true lack of records from these regions. Temporally, the Oligocene, Eocene and Paleocene are undersampled compared to the Quaternary and Pliocene, as shown in Figures 5 and 6.

**Figure 6.**
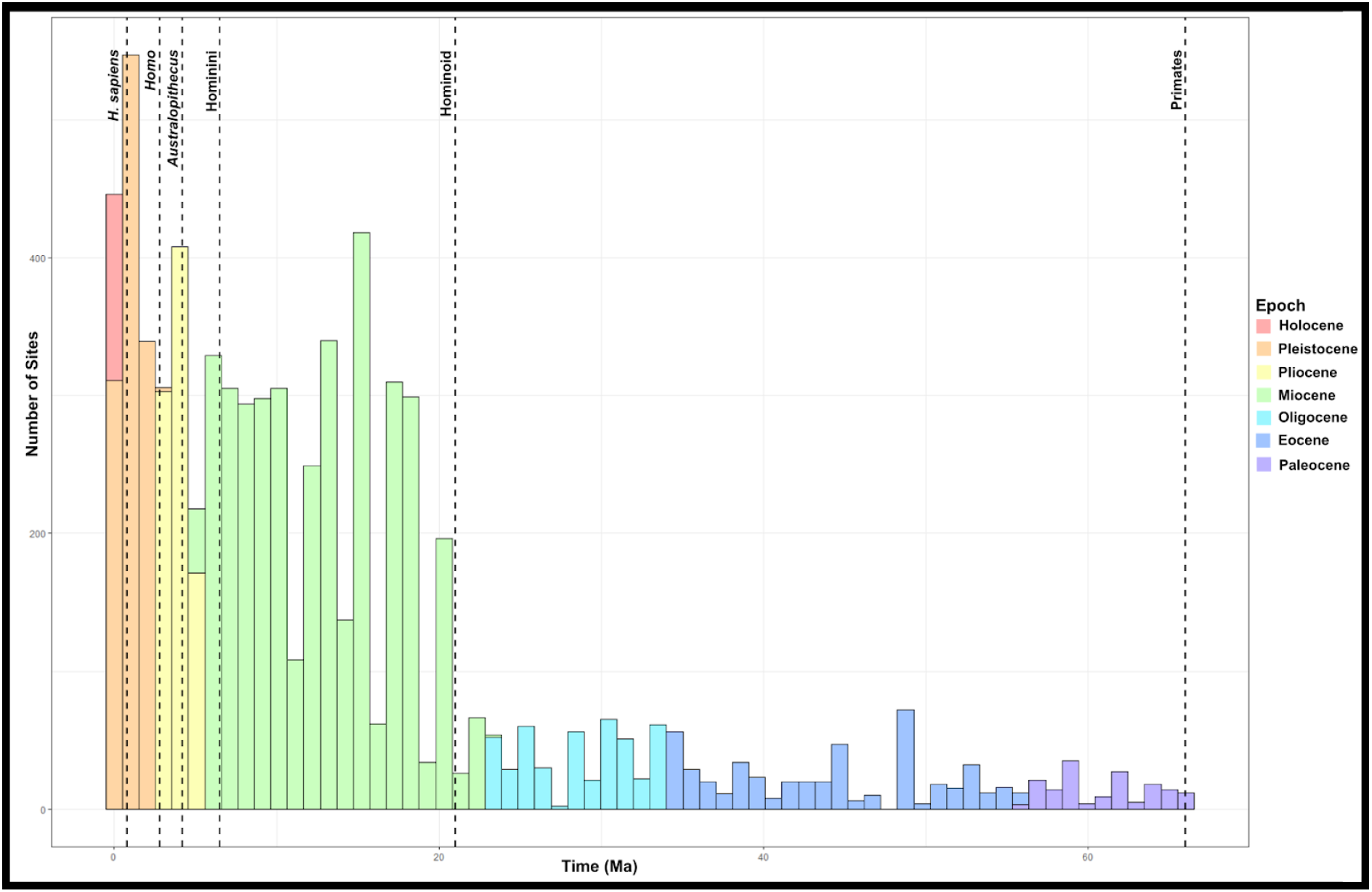
Histogram of number of global fossil mammal localities throughout the Cenozoic by 1 million year time bins. Dotted lines represent first appearance dates for clades of primates including *Homo sapiens*.

It is conceivable that what appears in this analysis as anthropocentric sampling bias is simply an artefact of sediment availability. Specifically, that the locations in the NOW database are a reflection of what is available in the fossil record, and what appears as an oversampling of localities and time slices at which hominins occurred in actuality reflects where and when sediment was abundantly deposited. We suggest that this is unlikely and that sampling is geographically biased in favour of spaces where hominins have been found, while areas in which hominins have not been found are undersampled (Fig. 4). For example, south-western Europe is comparatively basin-poor but has abundant fossil localities. Western Africa, in comparison, is basin-rich and has barely been sampled. Of the 16 nations defined by the United Nations as western Africa (https://unowa.unmissions.org/map) only Mali (2 sites), Niger (1 site) and Senegal (2 sites) are sampled, containing 5 fossil localities. Within these 5 sites, only Tobène (Early Pliocene, Senegal) contains more than 3 recorded taxa (Lihoreau et al., 2021).

Even within well sampled regions, such as the Eastern African Rift System (EARS), this anthropocentric bias is problematic. Recently, Barr & Wood (2024) have shown that the fossil record of the EARS is not representative of the range of environments occupied by most extant rift-dwelling mammals. In particular they (Barr & Wood, 2024) show that EARS represents arid, grassy and shrubby environments the best, while not representing the more wooded and wet environments in a species’ range. Therefore, it is highly unlikely that the eastern African record (which mostly comes from the EARS, Fig. 4) is a truly representative sample of hominin environments in Africa (Barr & Wood, 2024); and it is likely that environments with high precipitation and tree cover are amongst those which are not represented by the current African sample.

Our analyses do not account for whether the lower sampling reflects lower sediment availability in each time slice. Though we did not have suitable data to analyse this question, the ‘pull of the recent’ is a well documented phenomenon affecting the fossil record (Jones et al., 2021; Raup, 1972, 1979; Sepkoski, 1981). We did therefore anticipate that there would be an effect of decreasing sediment availability moving back through time. However, here we discuss only the Cenozoic fossil record, which in terms of geologic time, is ‘recent’. As such, we would not expect the reduction in fossil coverage due to the pull of the recent to give as marked a pattern as is seen in our results, particularly from the Early Eocene onwards (Sahney & Benton, 2017). Moreover, if the reduction in fossil sampling as we move back in time was only due to reducing sediment availability we would expect a more consistent reduction, rather than the very sudden temporal shift in number of sites we see in Figures 5 and 6. Therefore, we think it is likely that pre-hominoid time slices are underrepresented in the database, and that in many cases, particularly in specific geographical regions, the sediment is available but not exploited.

It is further important to discuss whether this result is an artefact resulting from the fact that hominins are present everywhere. However, we see here that in the fossil record (as recorded in the NOW database) this is not true. While modern humans are now present globally, altering most terrestrial biomes (Albuquerque et al., 2018), this is not true of the fossil record prior to the end of the Pleistocene. Non-sapient, human fossils are not present in many regions; including Australia, South America and Western Africa, which we have identified as undersampled (Figs. 2 and 3). In some regions, we can consider other biases which contribute to this effect. For example, western Africa is an undersampled region, where hominin fossils have not been discovered. It has been recognised (Almécija et al., 2021; Blinkhorn et al., 2022; Scerri et al., 2022) that this is a region in which hominin fossils, of great importance to our understanding of human evolution, may be located. Undersampling of this region is therefore likely not due to anthropocentrism, but due to geopolitical and socioeconomic factors, which prevent palaeo-scientists from prospecting in the region (Amano & Sutherland, 2013b; Hopkins et al., 2018). In addition to these limiting factors there are additional taphonomic limitations in much of western and central Africa. Much of the region has long been covered by rainforest, a poor preservational environment (Antonio Rosas & Saladie, 2022; Behrensmeyer et al., 2000). As such, even if potentially fossiliferous sediments have been deposited there the climatic and environmental conditions may not have been favourable for preservation (Behrensmeyer et al., 2000). However, limited work has been done to prospect in this region due to this presumption (Antonio Rosas & Saladie, 2022), again displaying neglect of the region.

An additional non-anthropocentric limiting factor to sampling of potentially fossiliferous sediments is accessibility. Ecological and geographic factors may prevent access to regions where fossiliferous sediments may have been deposited, or limit the visibility of sedimentary outcrops. For example in northern Europe (which we have highlighted as being poorly sampled) glacial erosion and heavy forest cover. As shown in Figure 4, the sediment deposition in northern Europe is already modest - sedimentary availability has been reduced post-deposition by glacial erosion during the ice ages (Helmens, 2014; Sejrup et al., 2005). Exacerbating this, heavy forest cover in the region (Verkerk et al., 2019) reduces visibility and access to any potentially fossiliferous sediment remaining. In northern Europe, recent finds have highlighted that despite the challenges of erosion, a Neogene fossil record is present and we consider this is likely to expand with further sampling effort (Saarinen & Salonen, 2024). In other regions, issues of accessibility, relating to heavy vegetation cover, topography or other landscape features, as well as political factors may exacerbate other factors that deter fossil prospecting in the region.

Simultaneously, there are issues of colonialism and the global power imbalance in palaeontology (Raja et al., 2022) which prevent some undersampled regions being sampled by ‘homegrown’ paleo-scientists (Cisneros et al., 2022). These cultural and social biases can also enhance anthropocentric sampling bias. The high concentration or fossil localities in southern and western Europe could be considered as a by-product of anthropocentric nationalism (Goodrum & Tech, 2009; Sautman, 2001) and specifically, the desire for the most important fossils to understand human evolution to be from ‘your’ country. One example of this is the Piltdown man hoax, in which British palaeo-scientists were so motivated to have Britain be central to the human evolution narrative that an unknown individual created a fake ‘missing link’ fossil (Goodrum & Tech, 2009). This story provides an extreme example of how anthropocentrism can corrupt the fossil record. While the fervour which drove the early 20th century race for human fossils has reduced, individuals can still progress their career and achieve fame by discovering human fossils providing motivation to prospect in hominin bearing locations, and promoting anthropocentric sampling.

It remains to consider the more insidious and ambiguous limitations anthropocentrism has, and continues to introduce to the fossil record. Narrowly, if we take the anthropocentric view of anthropocentrism, we can consider how the human fossil record itself is limited by this form of sampling bias. If we are only prospecting for hominin fossils, where we already know hominins are found, then we are potentially missing hominin sites which would expand our knowledge of our lineage and challenge our pre-formed hypothesis (Wood & Smith, 2022). For example, the evolution of hominins has long been connected with the arid savannah habitats of East Africa (Barr & Wood, 2024; Bender et al., 2012; Foister et al., 2023; Scerri et al., 2022), as this is where hominins are known to have existed. How would this hypothesis be challenged if contemporary hominin species were found in the forests of Western Africa (Scerri et al., 2022)? Applying this thinking broadly to the fossil record as a whole, we propose that we are almost certainly neglecting many mysteries of the fossil record by focusing on and sampling ecosystems which we believe to have affected hominin evolution.

This again is an issue which is affected by other sociopolitical issues and more practical ones, such as financial inequality. Ultimately, it is always going to be more economical to organise excavations close to home, particularly when this involves the training of students in palaeosciences. Researchers from the global south face specific challenges, such as visa costs, delays and bureaucracy, which often limit their ability to travel and work outside of their home countries (Chugh & Joseph, 2024). Raja et al., (2022), demonstrated that there is a European and North American monopoly on palaeontological knowledge, as a consequence of both resources and a culture of palaeontological research. This may in part explain the outlier of North America in our results, as the concentration of paleontological expertise in North America increases the likelihood of excavation in the region. There is an additional gender paradigm involved in travelling for fieldwork, as for female fieldworkers it has often been safer and more plausible to excavate locally (Burek & Kölbl-Ebert, 2007). This gendered dimension is one aspect of the historical traditions in fieldwork which have shaped the field. The fossil record has been growing since the work of Cuvier in the 19th century (and even before with ancient records of fossil discoveries), over the last two centuries of ‘formal’ palaeontology, the culture of research and practices have evolved and will continue to as the field develops. Perhaps anthropocentrism is a maladaptive strategy within this culture and practices, which it is time to move past.

We have applied the NOW database to evaluate the question of anthropocentric biases. As a solely mammalian database, which is expertly curated with emphasis on accurate stratigraphy and taxonomy, it tends to contain fewer demonstrable errors than other alternatives like the Paleobiology Database (Prothero, 2015; Žliobaitė & Fortelius, 2022). The NOW database is however constrained by its history. Originally, upon its 1998 creation the database was devised as a record of the ‘Neogene of the Old World’, but it has since expanded beyond this original concept, officially being renamed in 2012 to reflect it as a database of the ‘New and Old Worlds’(Žliobaitė et al., 2023). This history is a large part of why some areas, like western Eurasia, contain so many records. However, the database is continuing to grow, and particular effort is being diverted towards filling in some of the gaps identified here. Because it is a living database, it offers an opportunity to be able to correct the biases we have identified.

Ultimately, the problem of anthropocentric sampling bias reminds us of the importance of asking ourselves - ‘what questions can palaeontological data answer, and which are the most important?’ (Kiessling et al., 2019). We must consider if we want to understand the biotic world for how it functions intrinsically or only in how it functions in relation to human species. To investigate and limit the current global mass extinction event we must understand global ecosystems intrinsically and not only in relation to humans. Anthropocentric sampling bias severely limits the ability to do this. Moreover, if we wish to prepare for the climate change which will be experienced in the coming decades, centuries and millennia we need data about time periods with similar climates (Tierney et al., 2020). We identified the Eocene as an undersampled time slice (Figs. 5 and 6), which is precisely one of the time periods which may prove analogous to future Earth conditions (Tierney et al., 2020). The Miocene, while better sampled than the Eocene, is inconsistently sampled when we consider the time coverage of the fossil sites we have for this period (Fig 4b). Further sampling from these two epochs should be a high priority for future fieldwork.

## Conclusions

Ultimately, despite its biases and flaws the fossil record is the best tool we have in palaeosciences (Benson et al., 2021). This is exactly why it is so important to constantly expand our knowledge of the fossil record and interpretation of it. Towards this end, here we have analysed the sampling biases which anthropocentrism has and is introducing in the fossil record.

Our results are consistent with the idea that anthropocentric sampling introduces bias to the fossil record, specifically that fossil locality density is higher in time slices when hominins are found, and in known hominin-bearing locations. Therefore, in order to improve the quality of the fossil record and improve the ability of paleosciences to contribute to answering the most pressing questions facing the global scientific community, it is necessary to stop essentializing the narrative of human evolution in paleoscience, to reduce bias in sampling of fossil localities.

We acknowledge that this is a complex and difficult issue to address, which intersects with other matters, such as the difficulties in obtaining funding, social appeal of research and other factors we did not delve into here. We hope however, that this paper can act to draw awareness (Monarrez et al., 2022) to anthropocentric bias, and begin to shift focus of palaeosciences away from narratives centred on human evolution.

## Acknowledgements

We are thankful to Juha Saarinen for his helpful comments on this manuscript. We also acknowledge and thank Indrė Žliobaitė for her analytical guidance. O.E. Wilson acknowledges funding from the Research Council of Finland grant NEPA: non-analogue ecosystems of the past (grant number 340775/34692).

## Interest Statement

The authors report there are no competing interests to declare.

## Data Availability

The data used in the work were obtained from open access databases (Eco-ISEA3H database and the NOW database). All R code used for these analyses is archived at: https://github.com/TIFFOISTER/anthropocentrism.

## Notes

https://github.com/TIFFOISTER/anthropocentrism

### Competing Interest Statement

The authors have declared no competing interest.

